# Christmas disease in a Hovawart family resembling human hemophilia B Leyden is caused by a single nucleotide deletion in a highly conserved transcription factor binding site of the *F9* gene promoter

**DOI:** 10.1101/486373

**Authors:** Bertram Brenig, Lilith Steingräber, Shuwen Shan, Fangzheng Xu, Marc Hirschfeld, Reiner Andag, Mirjam Spengeler, Elisabeth Dietschi, Reinhard Mischke, Tosso Leeb

## Abstract

Hemophilia B is a classical monogenic X-chromosomal recessively transmitted bleeding disorder caused by genetic variants within the coagulation factor IX gene (*F9*). Although hemophilia B has been described in 28 dog breeds and four mixed-breed dogs hitherto, it has not yet been reported in the Hovawart. Here we describe the identification of a Hovawart family transmitting typical signs of an X-linked bleeding disorder. Five males had been reported to suffer from recurrent hemorrhagic episodes, four of them had to be euthanized finally and one died due to severe blood loss. A blood sample of one of these males with only 2% of the normal concentration of plasma factor IX (FIX) together with samples of seven relatives including the mother and grandmother were provided for further analysis. Next generation sequencing of DNA of the mother and grandmother revealed a single nucleotide deletion in the *F9* promoter (NC_006621.3:g.109,501,492delC; CanFam3.1). Genotyping of the deletion in 1,298 dog specimens (81 different breeds) including 720 Hovawarts revealed that the mutant allele was only present in the aforementioned Hovawart family. The deletion is located 73 bp upstream of the *F9* start codon in the highly conserved overlapping DNA binding sites of hepatocyte nuclear factor 4α (HNF4α) and androgen receptor (AR). The deletion only abolishes binding of HNF4α as demonstrated by electrophoretic mobility shift assay (EMSA) using purified recombinant human HNF4α and a transient overexpression lysate of human AR with double-stranded DNA probes encompassing the mutant promoter region. Luciferase reporter assays using wild type and mutated promoter fragment constructs transfected into Hep G2 cells showed a 65.3% reduction in expression from the mutant promoter. The data presented here provide evidence that the deletion identified in the Hovawart family caused a rare type of hemophilia B resembling human hemophilia B Leyden.

**Author summary:** Hemophilia B is the rarer form of classical hemophilias resulting from the absence or residual activity of blood clotting factor IX. Due to its X-linked recessive inheritance normally only males are affected. In a disease subtype, termed hemophilia B Leyden, factor IX activities increase during puberty resulting in spontaneous improvement of bleeding symptoms or even clinical recovery. This surprising development-related alteration is caused by nucleotide variants in important developmental and hormone-responsive regulatory regions of the factor IX gene promoter interfering with transcription factor binding. Although hemophilia B has been reported in several dog breeds, subtypes resembling human hemophilia B Leyden were unknown hitherto. In addition, the single nucleotide deletion reported here in Hovawarts in the overlapping binding sites of transcription factor HNF4α and androgen receptor only affecting HNF4α binding, was unexpected. Although it is advisable to genotype females in the future to prevent a further spread of this subtype of the disorder, our findings also open up the possibility not to euthanize affected males inevitably but to treat until puberty if necessary.

## Introduction

Hemophilia B (Christmas disease) is a recessive X-linked bleeding disorder caused by genetic variants within the clotting factor IX gene (*F9*) resulting in the absence or insufficient levels of factor IX (FIX) in the blood [1]. In humans hemophilia B is also known as the “royal disease” as it has been transmitted into several European royal dynasties by Queen Victoria [2, 3]. Currently, 1,113 unique *F9* variants have been described in man [4]. The majority of pathogenic variants is located within exons (923) and intronic regions (137) of *F9*. Only 33 variants (2.96%) have been described in the 5’-UTR (28) and 3’-UTR (5) accounting for 2.52% and 0.45% of human pathogenic hemophilia B variants, respectively [4].

Although first reports about canine hemophilia B date back to the early 1960’s and also being the first disorder in dogs characterized on DNA level, data on hemophilia B cases in dogs remain rather scarce compared to humans [5-8]. For instance, in the Cairn Terrier colony of the Francis Owen Blood Research Laboratory (University of North Carolina at Chapel Hill) a G>A transition (NC_006621.3:g.109,532,018G>A) in exon 8 causing an amino acid exchange (NP_001003323.1:p.Gly418Glu) was detected resulting in a complete lack of circulating FIX in affected dogs [9]. Due to a complete deletion of *F9* in Labrador Retriever, production of FIX inhibitors was detected after transfusion of canine blood products [10]. In two unrelated Airedale Terrier breeds a large deletion of the entire 5’ region of *F9* extending to exon 6 and a 5 kb insertion disrupting exon 8 was described, respectively [11]. Similar to the hemophilia B in the Labrador Retriever in both breeds FIX inhibitors were produced. A mild hemophilia B in German Wirehaired Pointers was caused by a 1.5 kb Line-1 insertion in intron 5 of *F9* at position NC_006621.3:g.109,521,130 [12]. Until today, hemophilia B has been described in four mixed-breed dogs and 28 dog breeds, *e.g*. German Shepherd, Lhasa Apso, Labrador Retriever, Rhodesian Ridgeback, Airedale Terrier, Cairn Terrier, Maltese and German Wirehaird Pointer [9-17].

In the canine cases analysed so far on DNA level, mutations have been observed only in exons and introns of *F9*, whereas alterations of the *F9* promoter have not yet been reported. In humans promoter variants have been detected resulting in the so-called hemophilia B Leyden characterized by low levels of FIX until puberty, whereas after puberty FIX concentrations rise to almost normal levels [18-20]. Since the first description, the genetic background of human hemophilia B Leyden was elucidated by various studies identifying variants in different transcription factor binding sites in the *F9* promoter including androgen-responsive element (ARE), hepatocyte nuclear factor 4α (HNF4α), one cut homeobox (ONECUT1/2) and CCAAT/enhancing-binding protein α (C/EBPα) binding sites [21, 22]. A special variant is hemophilia B Brandenburg resulting from variants in the overlapping binding site of HNF4α and AR [23, 24]. Unlike the classical hemophilia B Leyden, in patients with these variants FIX levels cannot be restored by testosterone-driven AR activity and remain low after puberty with no clinical recovery [21, 24].

## Results and Discussion

Hemophilias are rare diseases in dogs and hence it was rather coincidental that a case in a Hovawart (3, Fig 1) was reported to us. With the reconstruction of the pedigree it was possible to trace back the disease to the female conductor 39 (Fig 1). In the studied family the hemophilia was transmitted to 19, 4 and 6. 19 had one litter with 3 hemophilic males (48, 51, 53). 4 and 6 had litters with 1 affected male 60 and 3, respectively. Although DNA samples of 48, 51, 53 and 60 were not available, blood parameters and medical reports about recurrent hemorrhagic episodes were provided (Table 1). These males had increased activated partial thromboplastin times (aPTT) of 47.8 sec (53) to 72.9 sec (60) indicative for defects of the intrinsic coagulation pathway and also reduced FIX concentrations in the blood as is normally the case in hemophilia B. The affected dog 3 presented only 2% of the standard FIX concentration. The female conductors 4 and 6 showed aPTT within, however, FIX concentrations slightly below the reference range. The clinical signs together with the blood coagulation parameters and X-chromosomal transmission supported the diagnosis of a hemophilia B. The definite clinical diagnosis prompted us to search for the molecular cause initially on DNA level. The canine *F9* gene is located on chromosome X (CFAX) between positions 109,501,341 (transcription start site) and 109,533,798 and has a length of 32,458 bp (NC_006621.3, CanFam3.1). Similar to other mammals, the canine *F9* gene harbours 8 exons with an open reading frame of 1,356 bp coding for 452 amino acids [25]. DNA of female conductors 4 and 6 were subjected to whole genome sequencing and aligned to the canine reference *F9* gene sequence. Surprisingly, only 6 sequence variants outside the coding regions of *F9* were identified (Table 2). Five variants were located in introns and were excluded as cause for hemophilia B in the Hovawarts. The remaining variant (deletion) was located in the promoter of *F9* 73 bp upstream of the start codon. As this deletion was located within a putative transcription factor binding site of hepatocyte nuclear factor 4α (HNF4α) and androgen receptor (AR) which had been shown in humans to be important for *F9* expression and mutated in hemophilia B Leyden and Brandenburg [23, 24], this position was analysed in more detail.

**Table 1.** Determination of hemophilia relevant blood parameters and medical reports

| ID<sup>a)</sup> | Sex | aPTT (s)<sup>b)</sup> | FIX (%)<sup>c)</sup> | Medical report |
| --- | --- | --- | --- | --- |
| 3 | m | n.d.<sup>d)</sup> | 2 | severe bleeding after chipping, blood transfusion, death caused by blood loss |
| 4 | f | 12.7 | 92 |  |
| 5 | f | 13 | 64 |  |
| 6 | f | 13 | 58 |  |
| 7 | m | 12.1 | 83 |  |
| 48<sup>e)</sup> | m | 49.6 | 3 | slight bleeding during second dentition, lameness/joint problems (age 4 months), several blood transfusions |
| 50 | m | 14.4 | n.d.<sup>d)</sup> |  |
| 51<sup>e)</sup> | m | 56.2 | 70 | umbilical hernia with internal bleeding after surgery, blood transfusion, minor bleeding episodes |
| 52 | m | 29.7 | 110 |  |
| 53<sup>e)</sup> | m | 47.8 | n.d.<sup>d)</sup> | recurrent slight bleeding, prolonged bleeding during second dentition, lameness/joint problems |
| 54 | m | 12.9 | 55 |  |
| 60<sup>e)</sup> | m | 72.9 | 5 | severe bleeding after first vaccination (age 8 weeks) |
| Control 1 | m | 11.8 | 83 | healthy unrelated Hovawart control |
| Control 2 | f | 11.6 | >100 | healthy unrelated Hovawart control |

a) Animal IDs refer to Fig 1; b) aPTT (s): reference range 10-13.1; c) FIX (%): FIX % of standard (reference range: 75-140%); d): n.d.: not determined; e) 48, 51, 53 and 60 had been euthanized.

**Fig 1.**
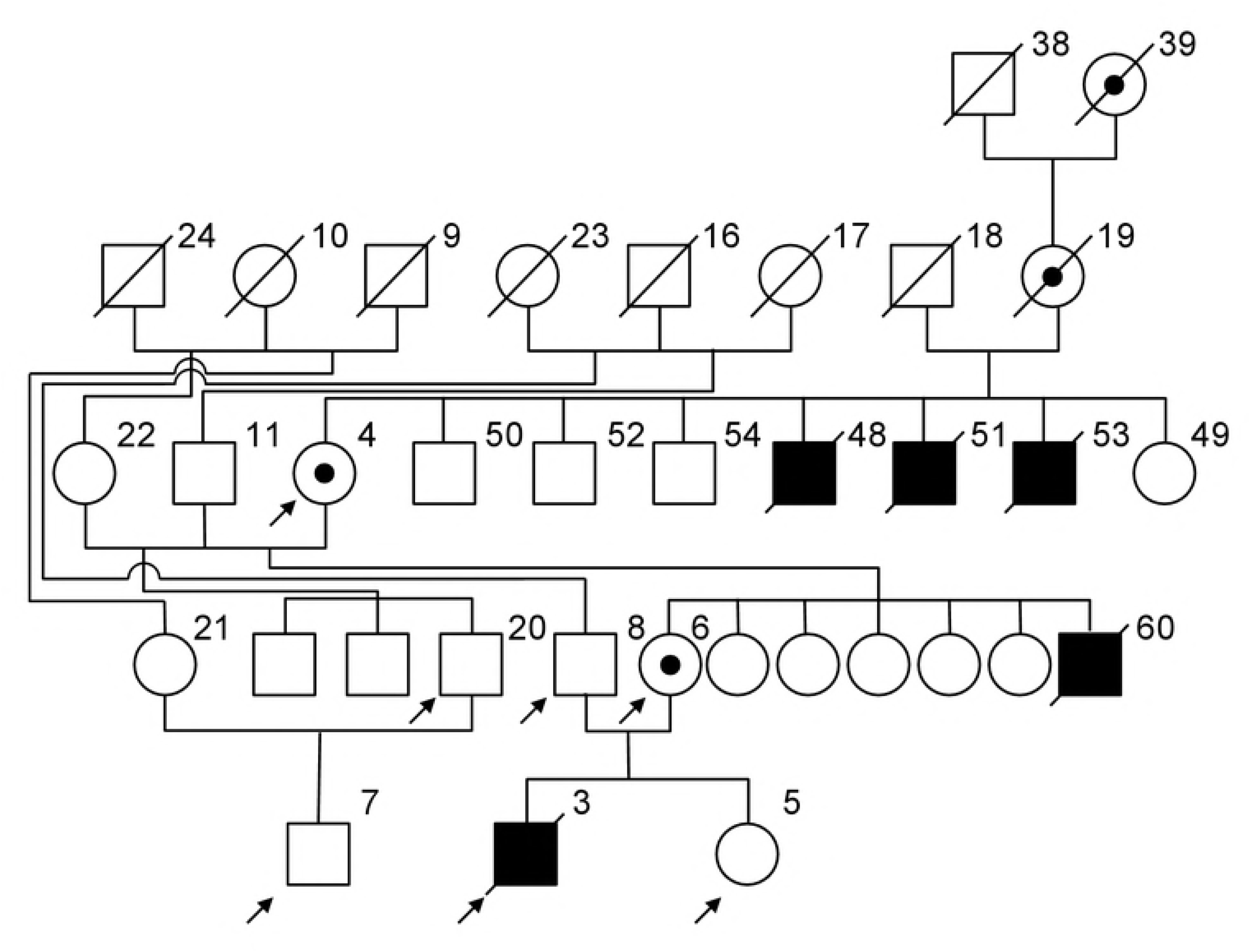
Pedigree section of the hemophilia B Leyden Hovawart family Pedigree symbols are according to the standardized human pedigree nomenclature [40]. Individuals are pseudonymized using internal IDs. DNA samples were available of individuals indicated with an arrow. For males 48, 51, 53 and 60 hemophilic signs (Table 1) have been reported and the dogs had to be euthanized after recurrent hemorrhages.

Figure 2 shows the segregation of the nucleotide deletion in the affected Hovawart family. The female conductors 4 and 6 were heterozygous, evident by the overlapping peaks with similar heights 5’ of the deletion position. The affected male 3 was hemizygous for the deleted allele whereas his sister 5 and cousin 7 were homozygous wild type. Genotyping of 1,298 dogs (including 83 different breeds, 720 unrelated Hovawarts, 12 Hovawart family members) demonstrated the occurrence of the deletion only among members of the affected Hovawart family (Table 3). To provide proof that the deletion represented the causative genetic variant and resulted in the low expression of *F9*, electrophoretic mobility shift and luciferase reporter assays were performed.

**Table 2.** DNA sequence variants in the canine *F9* gene determined by next generation sequencing of 4 and 6

| Position | Ref/Alt<sup>a)</sup> | Gene region | HGVS<sup>b)</sup> g. |
| --- | --- | --- | --- |
| X:109501492 | C/- | 5'-flanking region | NC_006621.3:g.109501492delC |
| X:109504229 | C/- | intron 1 | NC_006621.3:g.109504229delC |
| X:109505462 | -/AG | intron 1 | NC_006621.3:109505462_109505463insAG |
| X:109507446 | -/A | intron 2 | NC_006621.3:109507446_109507446insA |
| X:109510986 | G/A | intron 3 | NC_006621.3:g.109510986G>A |
| X:109524055 | A/G | intron 6 | NC_006621.3:g.109524055A>G |

a) Ref/Alt: Reference nucleotide/Alternate nucleotide; b) HGVS: Human Genome Variation Society (http://www.hgvs.org)

**Table 3.** *F9* genotype frequencies

| Genotype | Hovawart |  |  |  | Other breeds<sup>b)</sup> |
| --- | --- | --- | --- | --- | --- |
|  | HB<sup>a)</sup> affected<br>(n=1) | HB carrier<br>(n=2) | Control, related<br>(n=12) | Control,<br>unknown<br>relationship<br>(n= 720) | Controls<br>(n= 567) |
| C/C |  |  | 12 | 720 | 567 |
| C/- |  | 2 |  |  |  |
| -/- | 1 |  |  |  |  |

**Fig 2.**
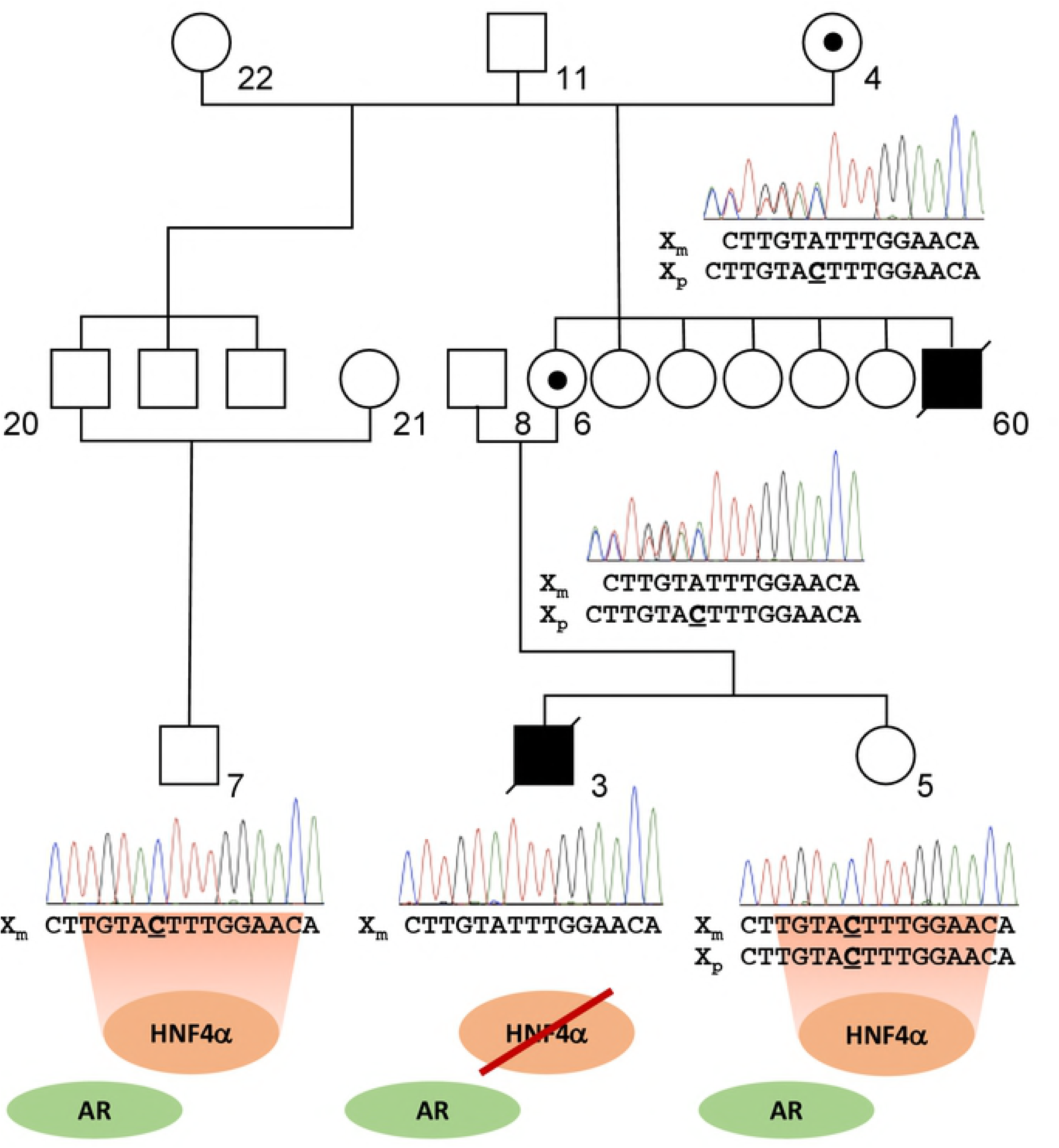
DNA sequence comparison of the mutatant hepatocyte nuclear factor 4α (HNF4α)/androgen receptor (AR) binding site in the promoter of canine *F9* in the hemophilic male (3) and relatives (4 grandmother, 5 sister, 6 mother, 7 cousin) The pedigree depicts a detail of the pedigree in Fig 1. Pseudonymized animal numbers also refer to Fig 1. Pedigree symbols are according to the standardized human pedigree nomenclature [40]. DNA sequences of heterozygous 4 and 6 (female conductors) show overlapping peaks with similar heights 5’ of the deletion position. X_m_: Maternal X-chromosome; X_p_: Paternal X-chromosome; HNF4α: Hepatocyte nuclear factor 4α binding site (consensus sequence: 5’-TGNACTTTG-3’) [21, 41]; AR: 3’-part of the androgen receptor binding site (consensus sequence: 5’-AGNACANNNTGTNCT-3’) [21, 41].

As shown in Fig 3 no binding of recombinant HNF4α to the mutated promoter region was detected. On the other hand, the AR lysate clearly showed binding to both fragments and hence the deletion seems not to influence AR binding to the androgen-responsive element in the canine *F9* promoter. This might be due to the fact that AR DNA-binding sites display an exceptional amount of sequence variation [26]. Although the C-deletion is located in the consensus TGTNCT-motif of class I AR-binding sites several alternative motives, *e.g*. TGTTTC in the stomatin-like protein 3 gene or TGTATC in the prostate-specific antigen gene enhancer III region, have been reported [26-28]. Therefore, it can be assumed that the affected males would have recovered from hemophilia during puberty. To analyse the effect of the promoter variant on *F9* expression, wild type and mutated promoter fragment luciferase constructs were transfected into Hep G2 cells. As shown in Fig 4 the mutated promoter fragment resulted in a statistically significant reduction of gene expression to approximately 34.6 % of the wild type promoter. The remaining activity of the mutated promoter is in agreement with the clinical findings of a residual FIX activity in the affected males (Table 1) and the results of the EMSA showing binding of AR in androgen-dependent promoter activation.

**Fig 3.**
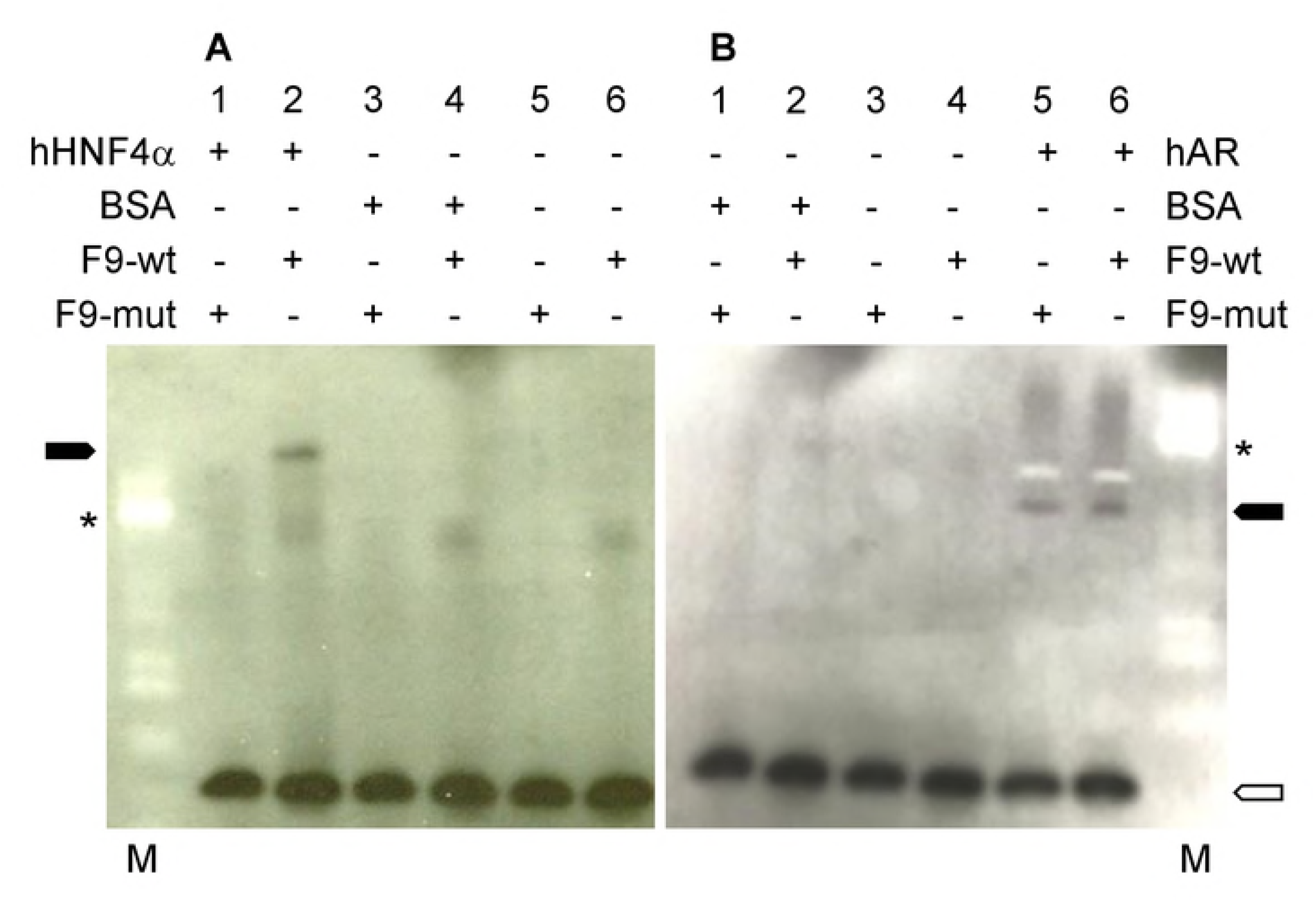
Analysis of HNF4α and AR binding of wild type and mutated *F9* promoter region using electrophoretic mobility shift assay Human HNF4α (A) and AR (B) were used to bind biotin-labelled wild type and mutated *F9* promoter fragments (F9-wt, F9-mut). Specific shifted bands (solid arrowheads) are detected in lane 2 (A) for HNF4α and lanes 5 and 6 (B) for AR. To test specificity, binding reactions were also performed using BSA (lanes 3 and 4 (A), lanes 1 and 2 (B)). In lanes 5 and 6 (A) and lanes 3 and 4 (B) no protein was added. Binding reactions were separated on 12% Tris-Glycine gels. X-ray films were cropped using GIMP 2.8.22. The 70 kDa protein marker band (PageRuler Prestained Protein Ladder, Fermentas) is indicated with an asterisk (lane M). The open arrowhead indicates unbound free DNA.

**Fig 4.**
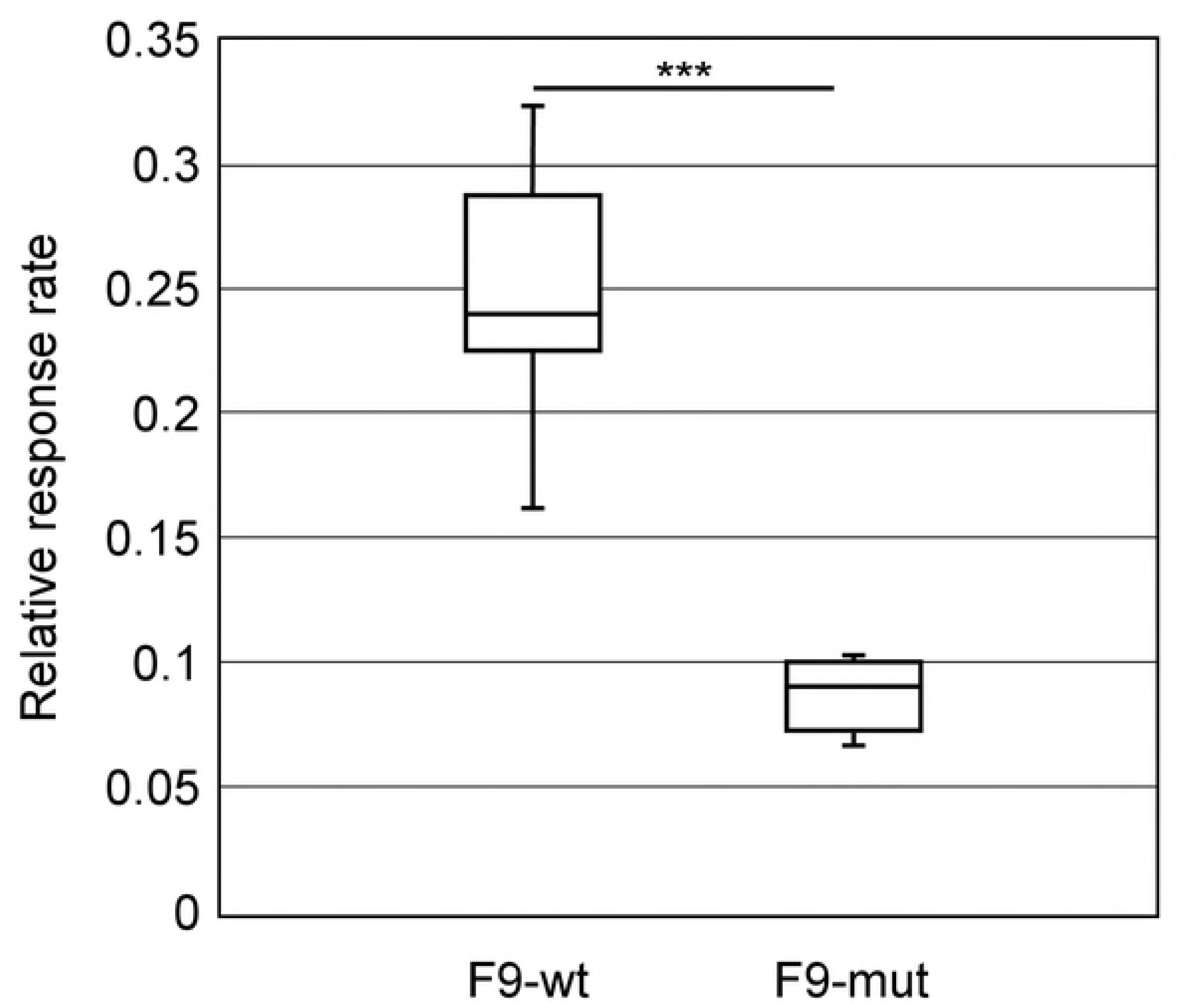
Dual-luciferase reporter analysis of *F9* promoter activities in Hep G2 cells Box and whisker plot showing the change of relative response ratios (RRR) between the wild type (F9-wt) and mutant promoter (F9-mut) gene constructs. The lines in the boxes represent the median. Whiskers indicate minimum and maximum RRR values. Values have been normalized as described above. Significance levels are indicated with asterisks (*p < 0.05, **p < 0.01, ***p < 0.001).Figure 1

In summary, we have identified and elucidated the causative genetic variant for hemophilia B Leyden in Hovawarts. This is the first report on a single nucleotide deletion within the binding sites of HNF4α and AR in the *F9* promoter causing hemophilia B Leyden in dogs. As the deletion only abolishes the binding of HNF4α, it can be assumed that male dogs will most likely recover during puberty as reported in humans [29-31]. However, to prevent any risk of a further propagation of the disorder genotyping of females is recommended in further breeding.

## Materials and Methods

### Ethical statement

The collection of dog blood and/or hair samples was done by local veterinarians. The collection of samples was approved by the Lower Saxony State Office for Consumer Protection and Food Safety (33.19-42502-05-15A506) according to §8a Abs. 1 Nr. 2 of the German Animal Protection Law (TierSchG).

### Animals and genomic DNA isolation

EDTA blood and/or hair samples were provided by different Hovawart and dog breeders with written owner consent. DNA was extracted from 30-50 hair roots using the QIAamp DNA Mini Kit (Qiagen, Hilden, Germany) according to the manufacturer’s instructions. A salting out procedure [32] was used for EDTA blood samples. Additional DNA samples deposited with the Institute of Veterinary Medicine were used as controls. All samples were pseudonymized using internal IDs.

### Coagulation assays and FIX activity measurement

APTT was measured coagulometrically using different commercially available activating reagents according to the manufacturer’s test instructions. To standardize measurement results performed in different laboratories, ratio values (aPTT patient/median aPTT of healthy dogs) were calculated and reported. FIX activity was measured coagulometrically using human FIX deficient plasma and a commercial human aPTT reagent for activation. Canine pooled plasmas were used as reference (activity defined as 100 %).

### Next generation sequencing (NGS) and genotyping

DNA of 4 and 6 was used for NGS on an Illumina HiSeq2500. A 450 bp library was prepared from genomic DNA with the NEBNext Ultra DNA Library Prep Kit for Illumina (New England Biolabs GmbH, Frankfurt, Germany) following the manufacturer’s instructions. Library quality was evaluated with Agilent2100 Bioanalyzer. Quality of fastq-files was analysed using FastQC 0.11.7 [33]. Total reads of 1,029,601,630 (4) and 1,000,503,256 (6) were obtained and mapped to the reference canine *F9* gene (NC_006621.3, region 109,501,341 to 109,533,798; CanFam3.1) using DNASTAR Lasergene Genomics Suite SeqMan NGen 15.2.0 (130) [34-36]. The following assembly options were used: mer-size 31 nt, min. match percentage 98, high layout stringency, min. aligned length 120 nt, min. layout length 50 nt, max. gap size 5 nt. Duplicate reads were combined and clonal reads removed. For 4 4,590 and for 6 4,982 consistent paired reads were assembled. The sample wise insert size metrics for high quality aligned reads was median pair distance 383.1 bp (SD 117.46 bp, min. distance 151 bp, max. distance 894 bp) for 4 and 383.9 bp (SD 103.55 bp, min. distance 153 bp, max. distance 882 bp) for 6.

Targeted genotyping of the promoter deletion was done by PCR amplification with primers cfa_F9_Ex1_F (5’-CCACTGAGGGAGATGGACAC-3’) and cfa_F9_Ex1_R (5’-CCCACATGCTGACGACTAGA-3’) resulting in a fragment of 328 bp (wild type) or 327 bp (deletion) spanning the variant position. The resulting PCR products were either directly sequenced on an ABI 3730 Genetic Analyzer (Thermo Fisher Scientific, Basel, Switzerland) or genotypes were alternatively determined by RFLP analysis after cleavage with *Rsa*I. In the wild type allele two fragments are generated with 52 bp and 276 bp while the allele with the deletion remained uncut.

### Electrophoretic mobility shift assay (EMSA)

For EMSA biotin-labelled double-stranded wild type (cfa_F9n_wt_Biotin: 5’-CAGAAGTAAATAC<u>AGCTCAACTTGTA</u>***<u>C</u>***<u>TTTGGAAC</u>AACTGGTCAACC-3’) and mutated (cfa_F9n_mut_Biotin: 5’-CCAGAAGTAAATAC<u>AGCTCAACTTGTATTTGGAAC</u>AACTGGTCAACC-3’) oligonucleotides were synthesized (Integrated DNA Technologies IDT, Leuven, Belgium) harbouring the overlapping HNF4α and AR binding sites (underlined). The position of the deleted C-nucleotide is indicated in bold and italics. Recombinant human HNF4α and human AR over-expression lysate were purchased from Origene Technologies Inc. (Rockville, USA). Binding reactions included 2 μL 10 × binding buffer (100 mM Tris, 10 mM EDTA, 1 M KCl, 60% v/v Glycerol (86% solution), 0.1 mg/ml BSA, 0.5% Triton X-100, 1mM DTT; pH 7.5), 2 μg poly(dI-dC) and 1.2 μg human HNF4α or 4 μg poly(dI-dC) and 5 μg human AR lysate. As negative controls 1 pmol duplex DNA oligos were incubated without protein or with 1 μg BSA. Binding reactions were pre-incubated for 20 min on ice followed by 1 hour at room temperature after adding 1 pmol biotin-end labelled double-stranded oligonucleotide probes. The mixtures were loaded onto 12% Tris-Glycine gels (Invitrogen, USA). After electrophoresis at 80 V for 90 min (HNF4α) or 2 hours (AR), gels were blotted onto PVDF membranes (GE Healthcare Life Sciences, Germany) using a wet blotter for 30 min at 100 V. Membranes were crosslinked at 120 mJ/cm^2^ using a commercial UV-light crosslinking instrument equipped with 254 nm bulbs for 1 minute. DNA detection was done employing the Chemiluminescent Nucleic Acid Detection Module Kit (Thermo Scientific, USA) with minor modifications, *i.e*. membranes were incubated for 1 min in the substrate working solution.

### Luciferase-Assay

For the luciferase assay the pGL3 Luciferase Reporter Vectors (pGL3-Basic, pGL3-Control) were used (Promega, Mannheim, Germany). The wild type *F9* promoter fragment (971 bp wild type) was generated by PCR using primers cfa9_HindIII_F_neu (5’-*CGTAGA CTTAGCACTGTTCA*<u>AAGCTT</u>CACACACACAGTT CTTAAAT-3’) and cfa9_HindIII_R_neu (5’-*ATGGCTAGCAACCGTCTAAG*<u>AAGCTT</u>AATTGTGCAAGGAGCAAGG-3’). The mutated *F9* promoter fragment (970 bp) was generated by PCR using primers cfa9_HindIII_F (5’-*ATCGTC*<u>AAGCTT</u>CACACACACAGTTCTTAAAT-3’) and cfa9_HindIII_R (5’-*CGTACG*<u>AAGCTT</u>AATTGTGCAAGGAGCAAGG-3’). For cloning into the *Hin*dIII restriction site of pGL3 primers were designed with an unspecific random 5’-tag (italics) followed by a *Hin*dIII restriction site (underlined). DNA of heterozygous female 6 served as template for amplification. Promoter fragment design was geared to an equivalence of the canine genomic situation choosing a respective distance between NC_006621.3:g.109,501,492delC and the luciferase start codon. Recombinant pGL3 vectors were used for transformation of *E. coli* XL1-Blue according to the manufacturer’s protocol. Plasmid DNA of 17 colonies of pGL3Basic+970bpinsertF9_MT and 37 colonies of pGL3Basic+971bpinsertF9_WT were isolated using Promega PureYield Plasmid Miniprep Kit (Promega, Mannheim, Germany) and sequenced for validation. A validated clone of each construct was incubated in LB-medium and plasmid DNA was isolated using Qiagen Plasmid Maxi Kit (Qiagen, Hilden, Germany).

For normalization *Renilla* luciferase activity was measured by co-transfecting phRL-TK(Int^−^) (Promega, Mannheim, Germany). Low expression levels of C/EBP in Hep G2 cells were complemented by co-transfection of a C/EBPα expression vector [22]. The carboxy-terminal triple FLAG human C/EBPα expression vector cloned in pcDNA3 was a kind gift of A. Leutz and E. Kowenz-Leutz (MDC, Berlin, Germany).

For analysis of promoter activity human hepatoma derived cell line Hep G2 (ATCC HB-8065) was cultivated in Roti-CELL DMEM High Glucose (Carl Roth GmbH, Karlsruhe, Germany) [37]. Constructs were transfected using Effectene Transfection Reagent (Qiagen, Hilden, Germany). Firefly and *Renilla* luciferase luminescence was measured using the Dual-Glo Luciferase Assay System (Promega, Mannheim, Germany) on a Tecan GENios Pro 96/384 Multifunction Microplate Reader (Tecan GmbH, Crailsheim, Germany) with the analysis software XFlour v4.64 after cell lysis with Passive Lysis 5X Buffer (Promega, Mannheim, Germany). Experiments were repeated 5-times with two measurements each. Background luminescence values were subtracted from raw luminescence values. *Renilla* luciferase activities were used for normalization [38]. Data are presented as relative response ratios [39]. To determine statistical significance Mann-Whitney *U* test was used. Values were considered statistically significant when *p < 0.05 (low), **p < 0.01 (medium) and ***p < 0.001 (high).

## Acknowledgments

The authors are grateful to S. Pach for expert technical assistance. L. Binder is thanked for support. The owners of Hovawarts who have provided blood samples are thanked for their generous support. A. Leutz and E. Kowenz-Leutz are thanked for providing the C/EBPα expression vector. S. Shan and F. Xu are supported with a fellowship by the Chinese Scholarship Council (CSC).

